# Structural Determinants for the Monomeric and Phospholipid-Transport P4B-ATPase via comprehensive *in silico* analysis of P4-ATPase structures

**DOI:** 10.1101/2025.07.01.662597

**Authors:** Kadambari Vijay Sai, Danny Farhat, Sai Anvithaa Sundar Rajan, Jyh-Yeuan Lee

**Author notes:** Correspondence addressed to: Jyh-Yeuan (Eric) Lee. **FOOTNOTE:** This report is part of a Special Issue commemorating the 68^th^ Annual Conference for Canadian Society of Molecular Biosciences: Protein Homeostasis – From Basic Science to Human Health, and the 4^th^ Biennial Proteostasis Researchers in Canada, Eh (PRinCE) Conference.

## Abstract

The P4-ATPase family of phospholipid flippases plays a critical role in maintenance of membrane asymmetry and cellular protein traffic and eukaryotic homeostasis. Several structures of these phospholipid flippases have been resolved, along with extensive biochemical characterization of the substrate transport properties. However, an essential subfamily of monomeric phospholipid flippases, the P4B-ATPases, remains to be characterized in depth. While P4B-ATPases appear to share similar lipid transport properties to their heterodimeric counterparts, the P4A-ATPases, the basis of their substrate translocation as monomers is currently unknown. In this study, we investigated the divergence of P4B-ATPases from other P-type ATPases using a structure-based analysis of sequence conservation. Our results showed conservative and non-conservative pockets and pathways in the P4B-ATPases near critical residues for the substrate transport pathway. P4B-ATPases also exhibit a unique interaction of an invariant proline near a conserved TM1-2 aromatic cluster, where dynamics simulation confirmed interactions with phospholipids and cholesterol by this conserved aromatic cluster. A critical TM6 residue was observed orienting into a conserved P4B-pathway whose function is currently unknown but a disease mutation hotspot, suggesting that P4B-ATPases exhibit novel transport and/or regulatory mechanisms. Our results provide a molecular framework for further studies on this essential subfamily of monomeric phospholipid flippases.

## INTRODUCTION

P-type ATPases constitute a large and essential family of membrane transporters in eukaryotic and prokaryotic organisms. This family of proteins, including phospholipid flipping P4-ATPases, hydrolyze ATP to generate a phosphorylated intermediate that can perform active substrate transport across the membrane (Palmgren 2023). Over the past few decades, structural studies of P-type ATPases have revealed the general molecular architecture of this transporter family, and different ligands have also been used to generate specific intermediates of the Post-Albers cycle (Stock et al. 2023). In the past few years, there has been a huge influx of P4-ATPase structures, particularly the P4A members (Sai and Lee 2024). These results significantly enhance our understanding of the structural and biochemical properties of their function. The P4B-ATPases, on the other hand, lack the same depth of biochemical and structural characterization.

P4-ATPases possess a similar overall architecture to the well-studied ion conducting P-type ATPases: a transmembrane domain with 10-12 transmembrane helices and three cytosolic domains including a nucleotide binding (N) domain, a phosphorylation (P) domain and an actuator (A) domain (Sai and Lee 2024). Among these, P4B-ATPases, which exhibit varying degrees of essentiality in eukaryotes, were predicted to have evolved earlier, whereas P4A and P4C-ATPases have developed the requirement for a *β*-subunit through evolution (Palmgren et al. 2019; Bai et al. 2021). P4B-ATPases include flippases such as Neo1 in *Saccharomyces cerevisiae*, TAT-5 in *Caenorhabditis elegans* and ATP9A and ATP9B in *Homo sapiens.* Despite their monomeric nature, P4B-ATPases carry out similar substrate translocation as P4A-ATPases. Interestingly, several residues of P4B-ATPases align structurally with other P-type ATPase groups, which is not seen in multiple sequence alignment (Bai et al. 2021). Although P4A- and P4B-ATPases share substrate specificity, the detailed transport mechanisms are well-defined for the P4A subclass, while but the pathways in the evolutionarily farther P4B subclass are not as extensively studied (López-Marqués et al. 2020).

This subfamily is considered essential in cell development., Deletion of Neo1, the sole P4B-ATPase in *S.cerevisiae* yeast, was shown lethal to cell survival (Hua and Graham 2003). Mammalian ATP9A, which has been implicated in neurodevelopmental disorders and hepatocellular carcinoma, is highly intolerant to missense mutations (Cordovado et al. 2025; T. Meng et al. 2023; Vogt et al. 2022; Mattioli et al. 2021; X. Wang et al. 2023; Yavas et al. 2026). Meanwhile, deletion of TAT-5, a *Caenorhabditis elegans* P4B-ATPase, was found to be embryonically lethal (Wehman et al. 2011; Beer et al. 2018). In addition, P4B-ATPases participate in intracellular trafficking, including ER-Golgi transport, endosomal recycling, and plasma membrane protein retrieval. For example, ATP9A regulates the recycling of Wntless, transferrin receptor, and GLUT1, while Neo1 interacts with trafficking adaptors (Ysl2-Mon2-Dop1 complex) to facilitate vesicle formation and COPI-dependant protein transport from the golgi (Barbosa et al. 2010; Hua and Graham 2003; McGough et al. 2018; Tanaka et al. 2016). Further, recent studies on these P4B-ATPases have highlighted the physiological roles of their flippase activities of PI4P by Neo1 or ATP9A, a function implicated in neomycin resistance phenotypes of yeast and humans and in their roles in intracellular traffic of viral proteins during infection (Yagi et al. 2025; Bhawik K. Jain et al. 2025). Pathogenic mutants of ATP9A have also been linked to reduced RAB GTPase activation, affecting its function in endosomal recycling, preventing neurite outgrowth and ultimately resulting in neurodevelopmental phenotypes (T. Meng et al. 2023). Given that some unique monomeric features of the P4B-ATPases have been studied in biochemical and cellular studies, a systematic characterization of their substrate transport pathway remains to be conducted to determine whether they possess conserved features that allow them to transport the same substrates despite being monomers. This will pave the way for further elucidation of their roles in protein traffic and disease phenotypes.

In this study, we performed a comprehensive sequence and structural analysis of P4B-ATPases against other P4-ATPases and P-type ATPase subfamilies and determined the conserved architectural features of the monomeric P4B transport pathway. Our results not only provide further insight into the structural evolution of P4-ATPases but will also serve as the basis for future mechanistic investigations of the understudied P4B-ATPase phospholipid transporters.

## RESULTS AND DISCUSSION

### Microenvironment for the Invariant Proline of P-type ATPase Transporters

We began analyzing the amino acid sequences of P4-ATPase phospholipid transporters from yeast, mammals, and plants, by incorporating features already known from P2-ATPase subfamily. Building on previous sequence analyses of P-type ATPases (Pedersen et al. 2012; Palmgren et al. 2019), we integrated our analysis with recent structural data of P4-ATPases by both cryo-EM and AlphaFold (Supp Table 1, 2). Mapping the conserved residues (Supp Table 3-5) onto previously resolved structures, we found that the transmembrane domains have several conserved residues in all P4-ATPases, with some only conserved in P4A- or P4B-ATPases and others conserved across all P-Type ATPases (Fig 1). First, the invariant proline of the TM4 PxxL motif (or PISL in P4-ATPases) exhibited a change in microenvironment between P4A- and P4B-ATPases (Fig 2A). These residues are critical for substrate selectivity, binding and stabilization as well as initiating dephosphorylation in P4A-ATPases (Mikkelsen et al. 2019; Nakanishi, Irie, et al. 2020; Coleman et al. 2012). This PISL motif is oriented toward the TM5 FFKY(N/S) motif and a conserved TM6 asparagine in all resolved P4A-ATPases, forming a Proline Pocket (Fig 2B). Mutations in this region have universally impaired substrate transport in P4-ATPases, including lethality in mutants of the essential Neo1 (Palmgren et al. 2019). However, in the P4B-ATPases, these regions are replaced by a TM5 VMHRG motif and a TM6 alanine (or serine) (Fig 2B). A recent cryo-EM structure of ATP9A showed that the arginine of the VMHRG motif coordinating the phosphatidylserine (PS) molecule along with water, in contrast to the lysine of the FFKY(N/S) of P4A-ATPases, which merely stabilizes the PS binding network (Abe et al. 2025). In our structural analysis, we also noted that the Proline Pocket appears to partially emerge in the substrate-bound state of ATP9A, although this was absent in the substrate bound state of Neo1 (Fig 2C). It remains to be seen whether these are differences that arose due to the different substrates and their different binding sites. The ATP9A and Drs2 structures also revealed that ATP9A^T882^ (and the equivalent Drs2^L1051^) were both found proximal to the substrate, but this residue is more conserved in P4B-ATPases. Since this residue exhibits greater conservation that ATP9A^S881^ in P4B-ATPases, it is possible that this residue is more critical for P4B-ATPase substrate transport. Ultimately, these fundamental differences suggest a divergent structural strategy for handling the same substrate in both P4A and P4B-ATPases, prompting us to examine the proline’s alternative microenvironment in P4B-ATPases.

**Figure 1.**
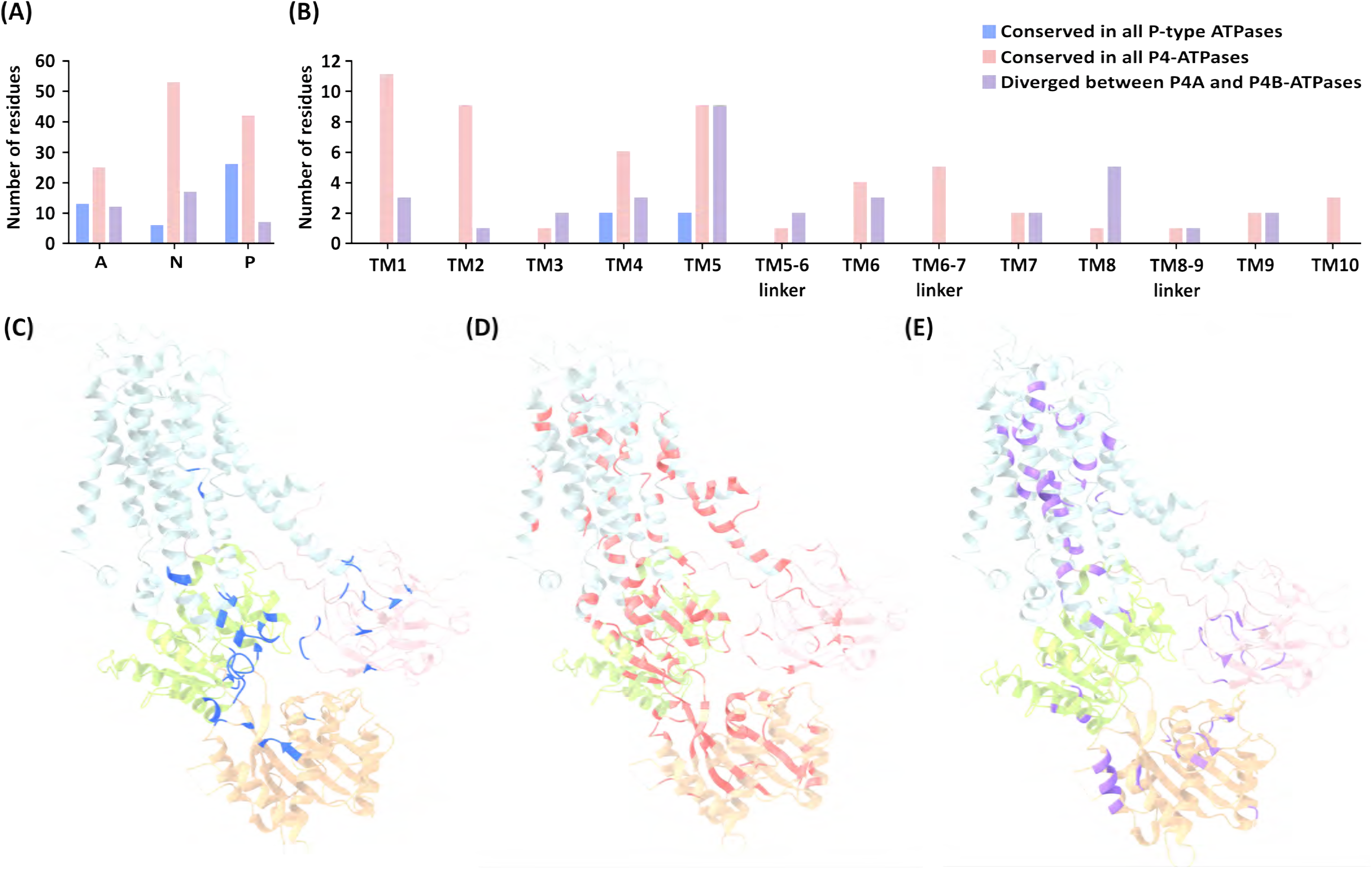
Distribution of conserved residues. Distribution of conserved residues in (A) cytosolic domains (B) transmembrane helices and their linkers, with the P-type (C), P4 (D) and P4A-B specific residues (E) mapped to the structure of Drs2 (PDB: 6PSX). Y-axis: number of conserved residues.

**Figure 2.**
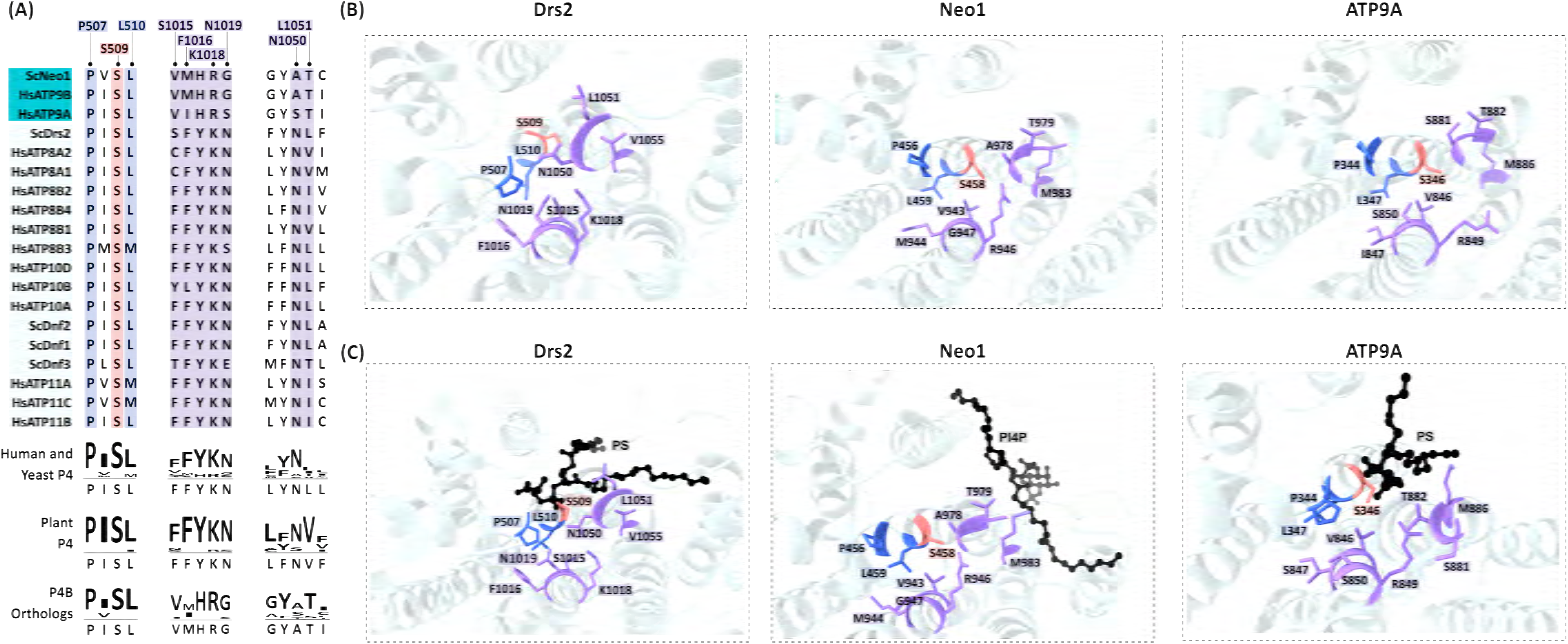
Adapting the Proline Microenvironment of P-type ATPases. (A) The PISL motif is a conserved motif in all P4-ATPases, which has a proline and leucine conserved in all P-type ATPases (blue) and a serine conserved in all P4-ATPases (coral). (B) Prior to substrate binding, this proline points toward a pocket formed by the P4A-specific motifs FFKYN and the TM6 Drs2^N1050^ (PDB: 6PSX), while P4B-ATPases such as Neo1 (PDB: 7RD6) and ATP9A (PDB: 9VDN) show this proline oriented completely away from this pocket. (C) Upon binding of substrates, the Drs2-PS (PDB: 7OH6) and Neo1-PI4P (PDB: 9BS1) bound structures exhibit similar orientations of the Proline Pocket, while ATP9A-PL (putative PS) (PDB: 9VDK) partially forms this pocket, although with the equivalent S881 oriented away from this pocket. Notably, the highly conserved residue in P4B-ATPases, ATP9A^T882^ orients into this pocket, nearer to the phospholipid substrate.

**Table 1.**
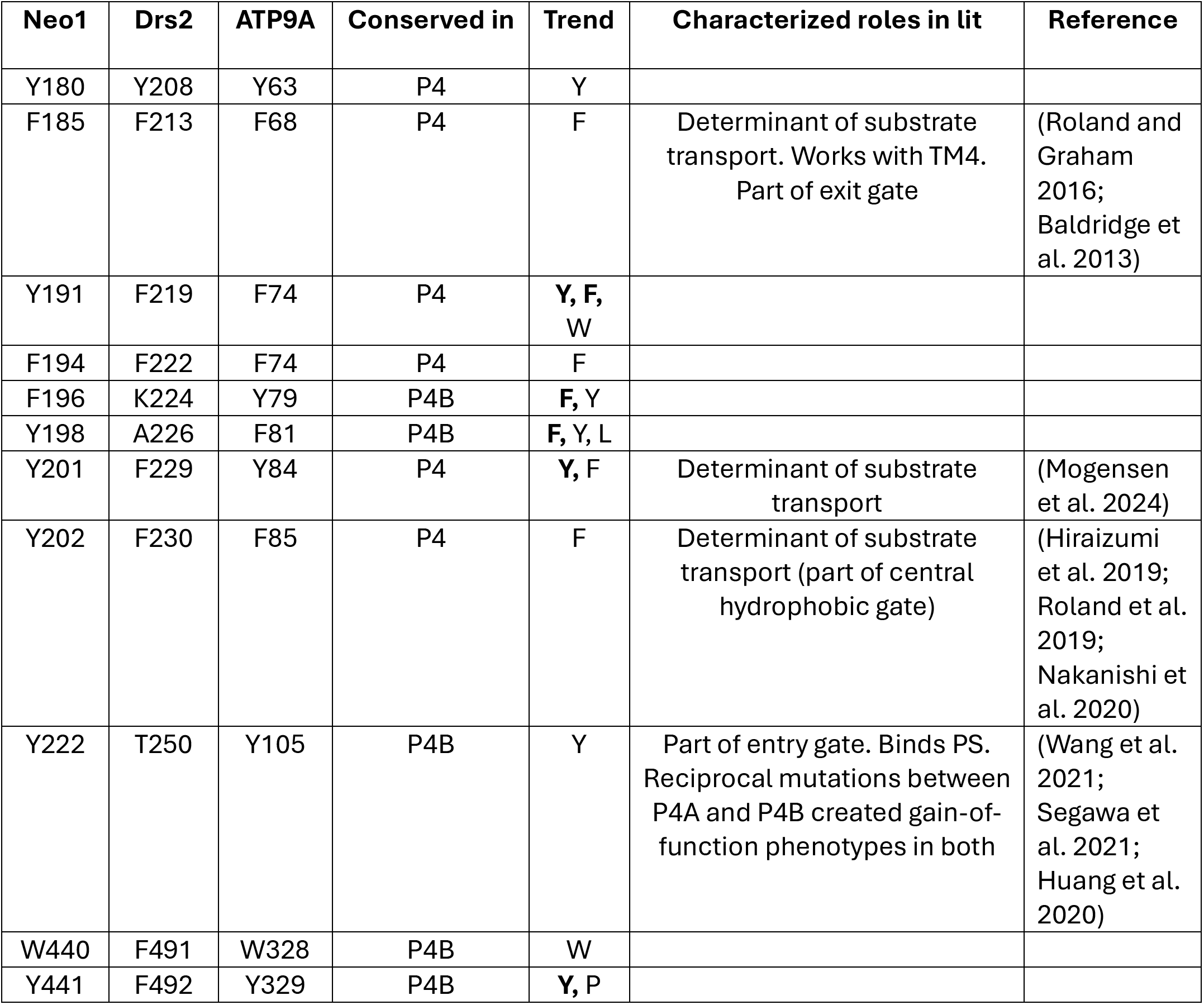
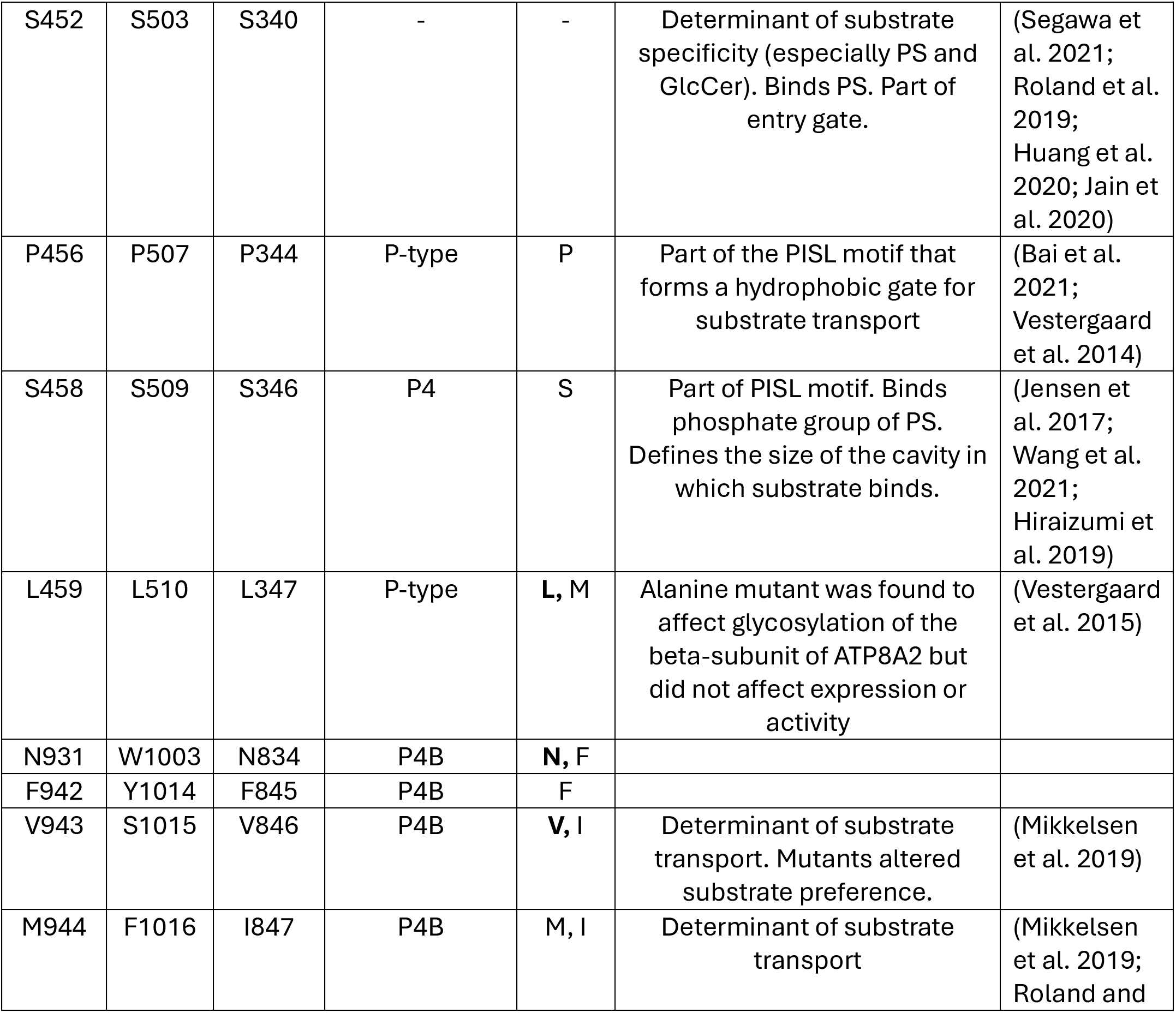
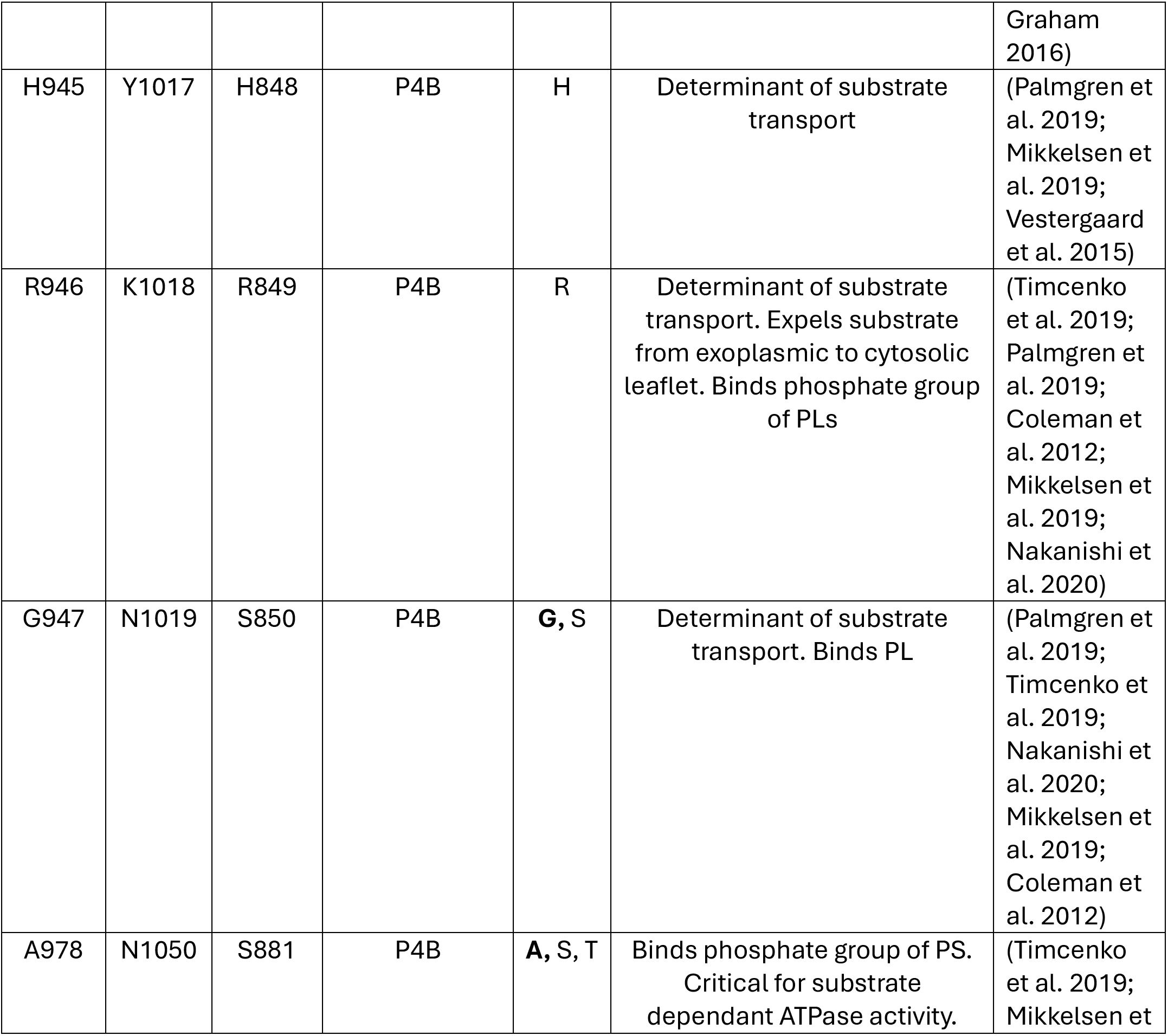
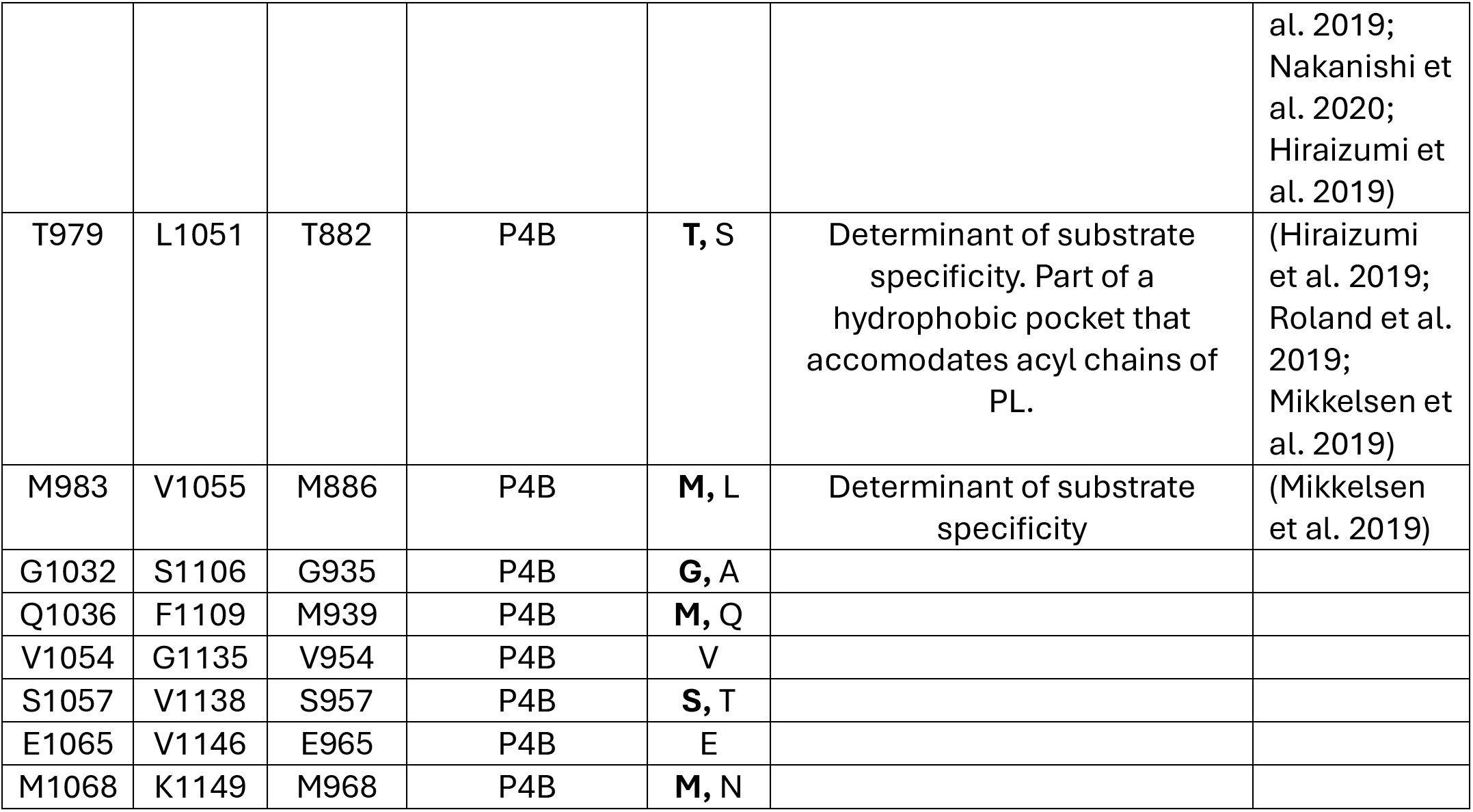
Identified motifs in this study. The residues highlighted in the study with their name and numbering in Neo1, Drs2 and ATP9A, listed in their order in the primary sequence of these proteins. Most of these residues are conserved in all P4-ATPases while some are throughout the P-type ATPase family or diverge between P4A and P4B. This table also highlights the characterized roles of these tables from previous structural, functional and cellular studies.

### Conserved Aromatic Aisle for Lipid-Transporter Interaction

Interestingly, the TM4 proline is oriented away from the VMHRG motif in P4B-ATPases but points to a previously unidentified TM1 Aromatic Aisle (Fig 3A), a conserved array of phenylalanines or tyrosines found in all P4-ATPases (Fig 3B). This proline headgroup is sandwiched between two residues of the Aromatic Aisle (Neo1^201YF^). Notably this Aromatic Aisle also contains two putative cholesterol sensing motifs ((L/V)-X_1-5_-Y-X_1-5_-(K/R) and (R/K)-X_1-5_-(F/Y/W)-X_1-5_-(L/V)). Previous studies suggested Neo1^F202^ is part of a hydrophobic gate that acts as a central barrier for alternating access to either side of the membrane and regulates substrate specificity, while Neo1^Y201^ contributes to PS transport (Mogensen et al. 2024; Vestergaard et al. 2014; Nakanishi, Irie, et al. 2020). It was speculated that some of these aromatic residues regulated transport of PS, phosphatidylehanolamine (PE) and glucosylceramide (GlcCer) on P4-ATPase members (Roland et al. 2019; Mogensen et al. 2024) but the aromatic group of these residues may contribute to interaction with lipids such as sterols, phosphoinositides or other glycolipids. For this, we performed coarse-grained molecular dynamic simulations on Neo1. Our results showed that palmitoyl-oleoyl phosphatidylinositol (POPI) and cholesterol molecules are stabilized and located along the Aromatic Aisle (Supp Fig 1, Supp Table 6). Conversely, established substrate lipids like palmitoyl-oleoyl phosphatidylserine (POPS) and palmitoyl-oleoyl phosphatidylethanolamine (POPE) exhibited short residence times in the transmembrane domain, suggesting that lipids like POPI and cholesterol remain bound to these transmembrane helices throughout the substrate transport cycle. Meanwhile, we observed that the proline was usually oriented between one or two phenylalanines of the TM6 FFKY(N/S) motif in P4A-ATPases (Fig 2C), suggesting the invariant proline and the aromatic residues required for positioning or recruitment of substrates is a shared principle across both P4A- and P4B-ATPases.

**Figure 3.**
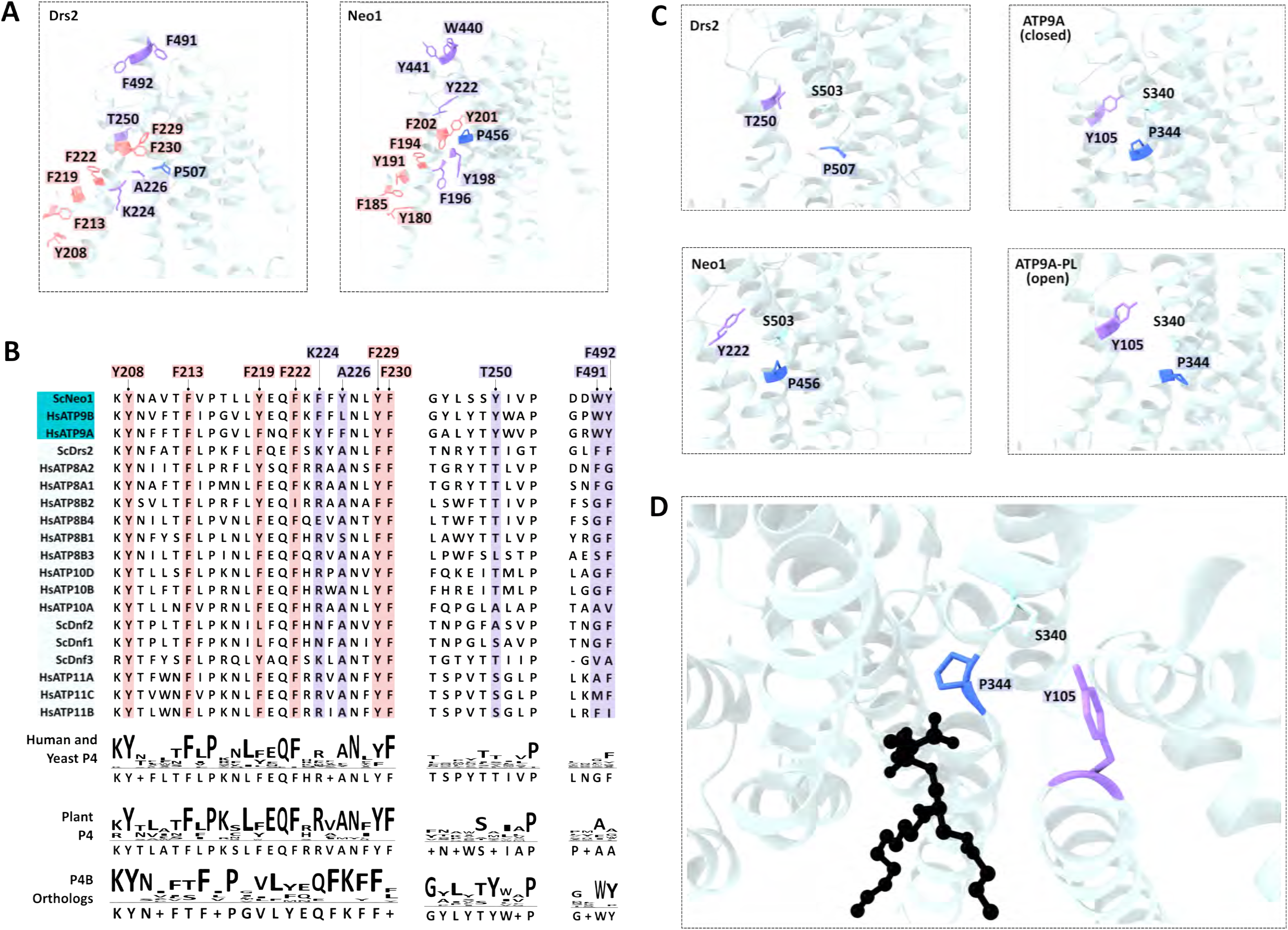
Aromatic Aisle and pathways of P4-ATPases. (A, B) In P4B-ATPases, the invariant proline of P-type ATPases orients toward a series of conserved aromatic residues conserved among P4-ATPases (coral). The P4B-proline is additionally surrounded by an aromatic cluster unique to the P4B-ATPase family (purple). (C) In this aromatic cluster, Neo1^Y222^ forms a pocket with the invariant proline unique to P4B-ATPases. (D) This pocket collapsed in the PL-bound state of ATP9A, where the proline is oriented toward the substrate instead of the pocket. The sequence numbers are based on the cryo-EM structure of an apo Neo1 (PDB ID: 7RD6).

### Conserved Featured Unique to P4B-ATPases

In addition to the Aromatic Aisle, the proline on P4B-ATPases is also surrounded by additional aromatic residues, Neo1^F196^, Neo1^Y198^ and Neo1^Y222^, which is not seen P4A-ATPases (Fig 3A). The functional importance of this architecture is underscored by mutagenenic studies. In P4A-ATPases, Neo1^F196^ (bovine ATP8A2^R54^) is part of a cytosolic electrostatic zipper that binds the substrate headgroup and thus prevents further substrate loading at the entry gate until the transport cycle is complete (Mogensen et al. 2024). In P4B-ATPases, however, it is not part of a similar ionic interaction; instead, it is instead closer to conserved residues such as the Y198 and other polar residues, which are critical for coordinating substrate transport and preference (Jensen et al. 2017; Nakanishi, Irie, et al. 2020; Roland and Graham 2016; Bhawik Kumar Jain et al. 2020; Palmgren et al. 2019). Meanwhile, Neo1^Y222^ is part of the entry gate and coordinates phospholipids in both P4A and P4B-ATPases. Strikingly, the Neo1^Y222S^ mutation is viable, whereas the reciprocal mutation in the P4A-ATPase Dnf1 (S243Y) confers novel PS transport ability. The lethality of a Neo1^Y222S/S452Q^ double mutant confirms that the pocket it forms near the proline is essential for P4B folding or function (Huang et al. 2020) (Fig 2C). A homologous pocket is present exclusively in the closed state of ATP9A, but is conspicuously absent in the open state bound to a phospholipid (Fig 2D), suggesting that this pocket is required before substrate binding and disappears upon substrate binding. In addition to this, Neo1^W440^ and Neo1^Y441^ also form parts of this P4B aromatic cluster that surrounds the proline, although at the distal extracellular end, potentially for sensing headgroups. Thus, P4B-ATPases have replaced the electrostatic gating mechanism of P4A-ATPases with a specialized aromatic architecture, creating a unique regulatory and substrate-coordination environment essential for their function.

Another interesting difference in the P4B proline microenvironment is that the critical TM6 ATP9A^S881^ (Drs2^N1050^) orients away from the proline and into a conserved pathway in the P4A-and P4B-ATPases, located between TM5, 7 and 8 (Fig 4A, B). Upon analysis of the pre-substrate-bound and substrate-bound structures of ATP9A, we found that the S881 orients toward R849 only in the latter, suggesting that it supports the interaction of PL with this critical residue (Fig 4C, E). In contrast to this, the equivalent P4A TM6 asparagine coordinates a water molecule which is eventually replaced by the phospholipid, which places it directly proximal to the substrate (Fig 2C). Mutagenesis of this residue drastically impairs function *in vitro* and causes neurological diseases such as Cerebellar ataxia, intellectual disability, and dysequilibrium syndrome 4 (CAMRQ4) (Alsahli et al. 2018; Hiraizumi et al. 2019; Dieudonné et al. 2023; Nakanishi, Irie, et al. 2020) (Supp Table 6). It is worth noting that such a difference in orientation of the equivalent Neo1^A978^ or Drs2^N1050^ was not observed (Fig 4C). While the unchanged nature of this residue in P4A-ATPases has been studied heavily, the interplay of this ATP9A^S881^ and other P4B-equivalents with R849 requires further characterization in the presence of phospholipid substrates. Apart from R849, several of the residues in the P4B Pathway that S881 orients toward at this stage remain a complete mystery. While studies in P4A-ATPases have shown that TM5 is critical for substrate transport and dephosphorylation by transmitting allosteric transmission of changes from the transmembrane domain to movement of the P-domain (Coleman et al. 2012; Nakanishi, Nishizawa, et al. 2020; Mikkelsen et al. 2019; Roland and Graham 2016), the roles of the highlighted residues in TM7 and 8 are unknown and require systematic characterization in both P4A and P4B-ATPases. Although there are no previous mutational studies on any of these TM7/8 residues, these TM helices have been identified as a mutational hotspot for diseases such as progressive familial intrahepatic cholestasis 1 (PFIC1), Parkinson’s Disease and other inborn genetic diseases, due to mutations in ATP8B1, ATP10B and ATP8A1 (Fig 4D, Supp Table 7). Structural and mutagenesis studies also implicate this interface in β-subunit (CDC50) binding and proper membrane trafficking, as chimeras or mutations in TM7-10 cause ER mislocalization and disrupted CDC50 glycosylation (Hiraizumi et al. 2019; Dieudonné et al. 2022; Vestergaard et al. 2015; Baldridge and Graham 2012; Stone et al. 2012). In addition to this, several lipids have been detected near these TM helices at interaction-permitting distances, though outside the pathway itself, including substrate lipids and regulatory lipids such as phosphoinositides, as well as other unidentified lipids (Hiraizumi et al. 2019; Dieudonné et al. 2023; Qian et al. 2025; Bai et al. 2020; Abe et al. 2025). This was also supported by our molecular dynamics simulations that revealed that POPI exhibited interactions at TMH7 and TMH10 (Supp Fig 1, Supp Table 6). Therefore, the conserved TM5/7/8 interface emerges as a critical multifunctional hub, integrating disease-associated mutations, subunit folding and stability, and the binding of both substrate and regulatory lipids.

**Figure 4.**
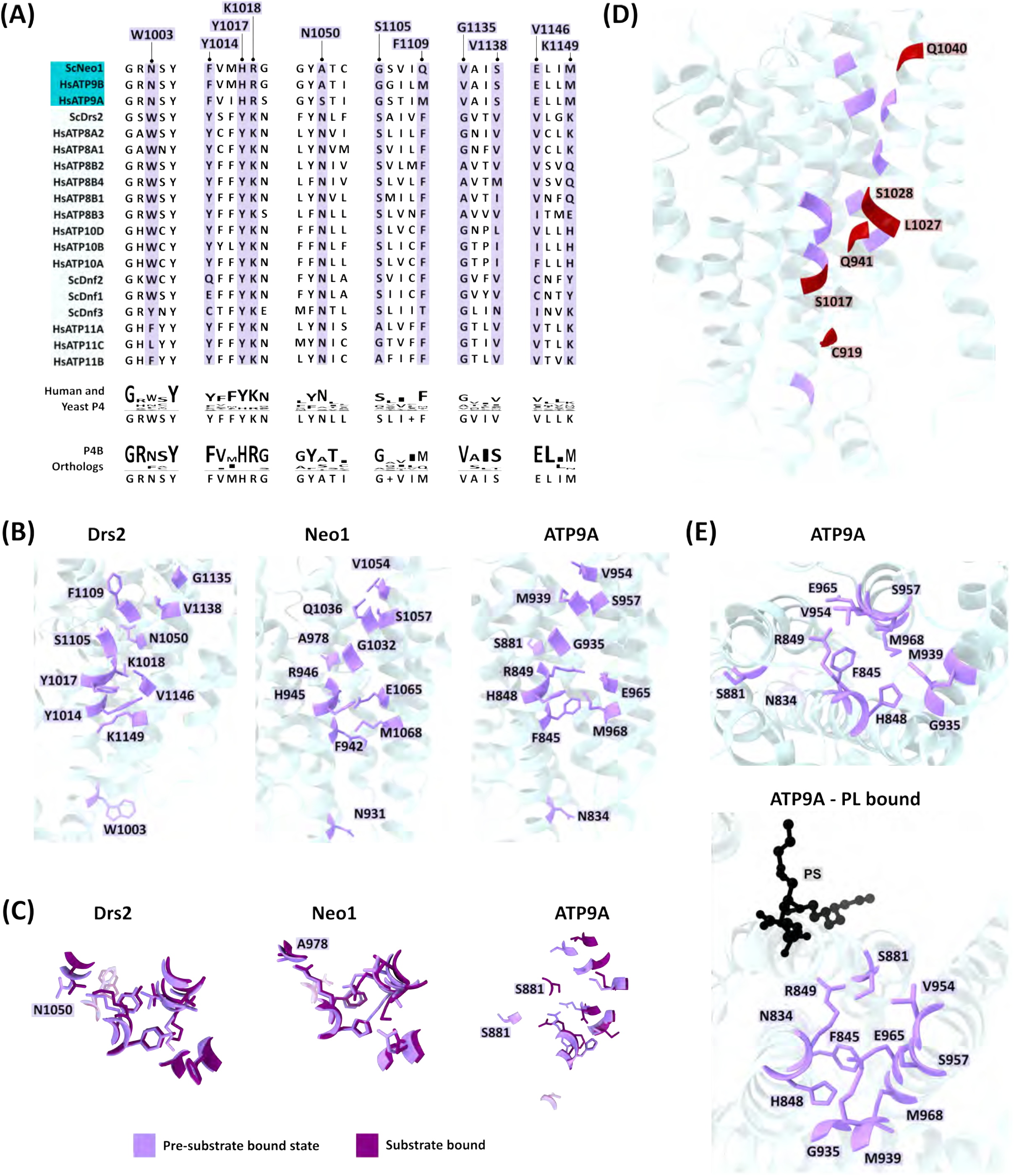
TM 5, 7 and 8 Pathway. (A, B) On the other side of the Proline Pocket lies a conserved pathway that diverges between P4A and P4B-ATPases, whose function is currently unknown, despite it being a disease mutational hotspot (red) as shown in (D). (C) The critical Drs2^N1050^ orients into this pathway independent of whether substrate is bound (dark purple) or not (lavender). Meanwhile the orientation of the P4B-equivalent appears to be more complicated, with Neo1-PI4P (dark purple) showing no difference from the Neo1-E2P (lavender), while ATP9A-PS appears to exhibit a change in position and orientation of the equivalent S881. (E) Despite this substrate induced change in the orientation of the equivalent ATP9A^S881^, inspection of the structures revealed that, upon substrate binding this residue is oriented away from the substrate and into this pathway.

## CONCLUDING REMARKS

Leveraging the several resolved structures of P4-ATPases and structure-prediction through AlphaFold, in combination with a sequence analysis-based approach, we have identified unique features of the P4B-subclass. This includes a novel microenvironment for a critical proline involved in substrate translocation in P4B-ATPases, a unique aromatic cluster involving an Aromatic Aisle in TM1 and another pathway between TM5, 7 and 8, which is a disease hotspot in P4A-ATPases. These require further investigation through exchange of P4A-B residues in biochemical studies, complemented by atomistic simulations of water molecules or phospholipids through the P4B-ATPase TM domain (Abe et al. 2025; Calianese et al. 2024; Y. Wang et al. 2021; Zhang et al. 2023; López-Martín et al. 2022; Dieudonné et al. 2023). This will shed light on how substitutions of critical motifs and residues near the substrate translocation pathway can support translocation of the same substrates as P4A-ATPases. Understanding how these two subfamilies diverge in their handling of the same substrates will inform future studies on their divergence of physiological roles. Eventually the results will also guide therapeutic strategies to rescue disorders such as PFIC which arise due to instability in P4A-ATPases. In addition to the divergence between P4A and P4B-ATPases, we also noticed a higher frequency of substitutions at several conserved residues in Dnf3, ATP8B3 and ALA1, suggesting functional impacts on these poorly understood substitutions. Moreover, specific substitutions were sometimes observed in ATP8B, ATP10 or ATP11 groups in humans, speculatively due to specific substrate transport or other functions. Conducting mutagenic studies on these residues, along with substrate-based phenotypic assays (such as papuamide A sensitivity for phosphatidylserine exposure) will inform how substrate specificity of different P4-ATPases works to regulate cellular processes. Ultimately, such studies will help understand why several P4-ATPases transport similar substrates and how they work together in their cellular systems.

In addition to the highly conserved cholesterol sensing motifs of the TM1 Aromatic Aisle, we identified other less conserved cholesterol sensing motifs and several conserved aromatic residues oriented toward the membrane in different P4-ATPases. This suggests that membrane composition could allosterically control different P4-ATPases differently, while all of them are also regulated by a common mechanism involving TM1 and conserved aromatic residues. Systematic mutagenesis of the snorkelling aromatic residues along the cholesterol-sending motifs may be undertaken in reconstituted systems of varying phospholipid-cholesterol ratios. Such studies will shed more light on how P4-ATPases, which are embedded in a pool of their own substrates, are regulated by membrane composition and fluidity against constitutive activation. Since phospholipids and their derivatives are critical pieces of cell signalling machinery for diverse cellular processes, such studies will help understand how these flippases integrate dynamic signals from the membrane to maintain membrane asymmetry and regulate protein traffic.

In conclusion, our comprehensive analysis reveals that while P4A- and P4B-ATPases share a superimposable core architecture and transport similar substrates, the monomeric P4B-subclass has evolved distinct structural solutions for key steps in lipid coordination and gating. The conserved motifs and P4B-specific variations identified here provide a framework for future mechanistic studies and structure-function analyses of these essential membrane transporters. By integrating these findings with biochemical and structural approaches, we can advance our understanding of how P4B-ATPases achieve lipid flipping and how their dysfunction contributes to human disease.

## MATERIALS AND METHODS

### Selection of P-type ATPases for the study

Yeast (*Sc*), plant (*At*) and human (*Hs*) P4-ATPases were used to understand sequence divergence between the heterodimeric (P4A, C) and monomeric P4-ATPases (P4B), while the human P-type ATPases were used to investigate phylogenetic relationships of P4B-ATPases to P-type ATPases. Only human proteins were selected for P-type ATPases due to a lack of a comprehensive list of all P-type ATPases in *Saccharomyces cerevisiae* and *Arabidopsis thaliana*. The P4B-ATPases used for this study were identified by manual curation of characterized members of this subfamily.

### Pairwise identity

Canonical sequences for P-type ATPases (Supplementary Table 1) were retrieved from UniProt (The UniProt Consortium 2025). Initial sequence analysis and pairwise identity for human P-type ATPases was conducted using ClustalOmega (Sievers and Higgins 2018). The resulting tree and identity matrix was visualized as a heatmap using GraphPad Prism.

### AlphaFold Structures

Although structures have been resolved for various P-type ATPases, there exist several discrepancies (experimental technique, resolved state, completeness). Consequently, AlphaFold structures (Abramson et al. 2024) (v4) were generated in the absence of any ligand for all proteins mentioned in Supplementary Table 2.

### Multiple sequence alignments

The AlphaFold structures were submitted to PROMALS3D (Pei et al. 2008) to generated structure-based multiple sequence alignments. Due to the 20-structure limit imposed by the server, sMSAs of the individual families were generated. Further, given the P4B-ATPase family’s evolutionary closeness to the P-type ATPases, the characterized P4B-ATPases were aligned to each P-type ATPase subfamily to determine conserved residues of P4A, B, P4 and P-type ATPases. Further, these P4B-ATPases were compared against human, yeast and plant P4B-ATPases to identify residues which differed between the monomeric and heterodimeric subfamilies. The individual subfamily sMSAs were analyzed using JalView (Waterhouse et al. 2009) to generate the Weblogos used in the figures.

### Identification of ligand-binding residues

Ligand-bearing structures of P-type ATPases were imported to ChimeraX (E. C. Meng et al. 2023). The residues in contact with the ligand were identified using default parameters in the Contacts tool (VDW overlap >/= 0.4 Å). The union of ligand binding sites of for each ligand for all P4 and other P-type ATPases was used to generate the heatmap of ligand binding sites.

### Molecular Dynamics Simulations

The structure of Neo1 (PDB ID: 7RD8) was retrieved from the protein databank and loaded into CHARMM-GUI (Jo et al. 2008). The protein was inserted into a heterogenous generic mammalian Golgi apparatus as described on the OPM server (Lomize et al. 2012). The membrane patch dimensions were 150×150Å. The system was then solvated with 13Å of excess water and ionized with 0.15M NaCl.

Coarse grained molecular dynamics simulations were performed using GROMACS2021.4 (Páll et al. 2020)with the MARTINI2.2 force field (Qi et al. 2015). The system first underwent a two-stage minimization protocol. Initially, a soft-core free-energy minimization was performed using the steepest descent algorithm for 5000 steps. This was followed by a second minimization for 100,000 steps or until the maximum force fell below 10 kJ mol⁻¹ nm⁻¹. Nonbonded interactions were treated with a reaction-field electrostatics model using a relative dielectric constant of ε = 15 and a cutoff of 1.1 nm. Van der Waals interactions employed a cutoff scheme with the Potential-shift-verlet modifier, also truncated at 1.1 nm. The system was then equilibrated for 100 ns using a time step of 20 fs, during which position restraints (force constant of 50 kJ mol⁻¹ nm⁻²) were maintained on the protein backbone beads. The temperature was coupled to a reference value of 310.15 K using the v-rescale thermostat (τₜ = 1.0 ps). Semi-isotropic pressure coupling was applied at 1 bar using the Berendsen barostat (τₚ = 4.0 ps) with a compressibility of 4.5 × 10⁻⁵ bar⁻¹. Production simulations ran for 6.25 μs per replicate using a 25fs time step. Temperature was controlled with the v-rescale thermostat (τₜ = 1.0 ps, T = 310.15 K), while pressure was maintained at 1 bar using the semi-isotropic Parrinello–Rahman barostat (τₚ = 12.0 ps, compressibility = 3 × 10⁻⁴ bar⁻¹). Each production trajectory underwent independent minimization and equilibration protocol.

The 2D lipid density heatmap was done using MDAnalysis (Gowers et al. 2016), Lipyphilic (Smith and Lorenz 2021), numpy (Harris et al. 2020). The lipid and cholesterol headgroups (PO4, OH) were used for the plotting. Further lipid residence time calculations were done using the PylipID package (Song et al. 2022).

## AUTHOR CONTRIBUTIONS

KVS and SASR carried out the initial multiple sequence alignment; KVS performed further structural and sequence analysis. DF performed the molecular dynamics simulation. KVS and JYL wrote the manuscript; JYL oversaw the project and acquired research fundings. All authors have read and agreed to the submitted version of the manuscript.

## ACKNOWLEDGEMENTS

We thank the Protein Biophysics Core Facilities for the computational support and Ms. Fatemeh Rezaei for insightful discussions regarding cholesterol sensing motifs. This work was initiated with the support of the University of Ottawa Virtual Research Opportunities Internships to KVS and SASR. DF is supported by a Canada Graduate Research Scholarship. The project was funded by the Natural Sciences and Engineering Research Council Discovery Grants (RGPIN 2018-04070 & RGPIN-2025-04349) to JYL.

---

**Supplementary Figure 1.**
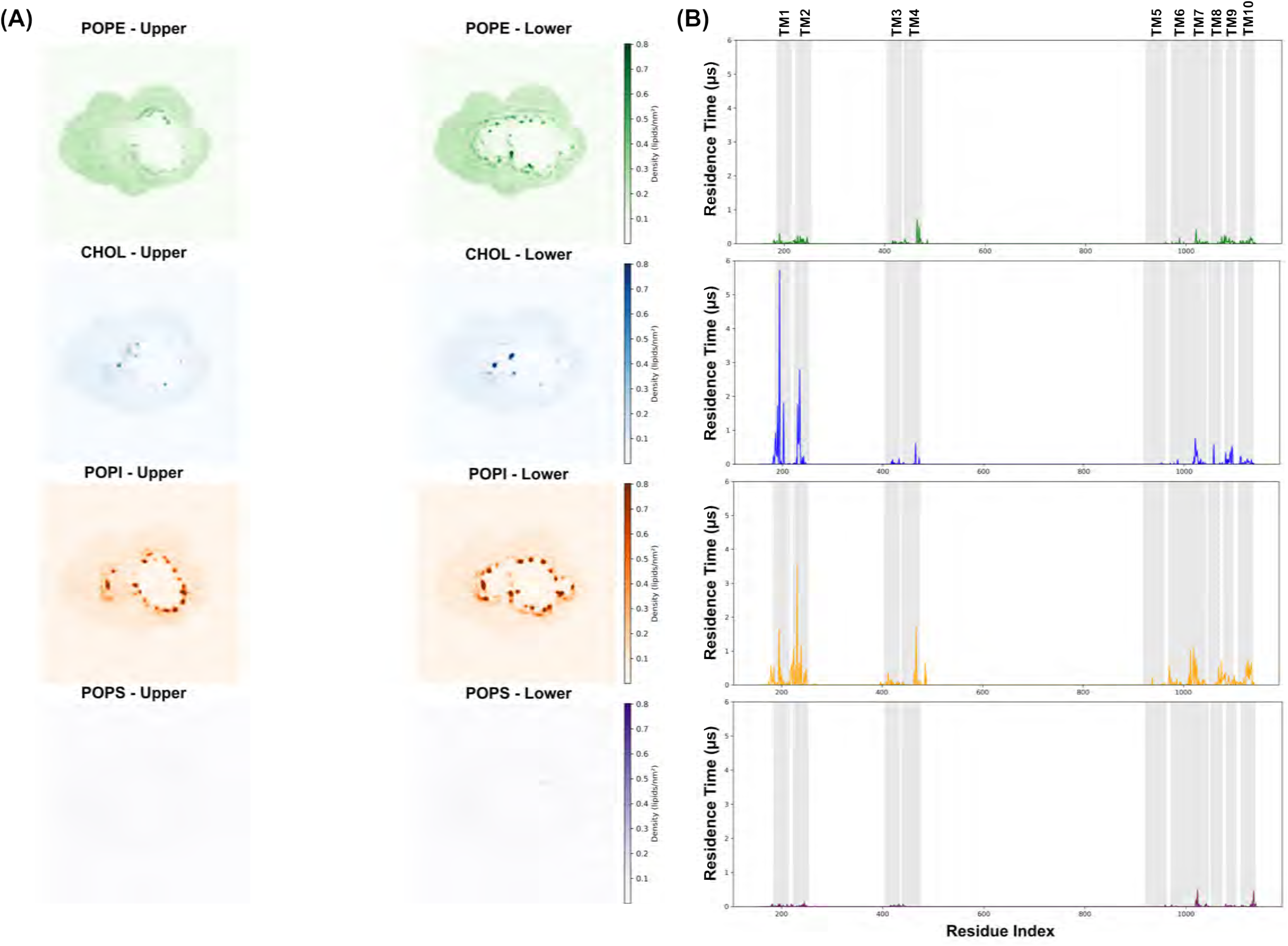
Lipid-protein interactions on Neo1 through coarse grained molecular dynamics simulations. (A) Heatmap of lipid density (POPE (green), Cholesterol (Blue), POPI (orange) and POPS (purple)) through three independent simulations on an Neo1 model (PDB ID: 7RD6). POPE and Cholesterol bind the protein in specific regions while POPI is less selective. (B) Residence time of each lipid to protein residues with the TMs highlighted in gray. Residence time was measure in microseconds as 1/Koff. POPE and POPS do not show strong residence times to the protein compared to Cholesterol and POPI which show strong association to TM1 and TM2 with some residual POPI binding to TM4 as well as both POPI and Cholesterol binding to TM7, 8, 10. TM sequence ranges on Neo1 (PDB ID: 7RD6): TM1 (185-215), TM2 (223-253), TM3 (405-434), TM4 (439-475), TM5 (920-961), TM6 (971-1001), TM7 (1003-1044), TM8 (1049-1072), TM9 (1080-1099), TM10 (1109-1137).

**Supplementary Table 1.** P-Type ATPases used in this study and their classification.

**Supplementary Table 2.** Structures of P-Type ATPases used in this study.

**Supplementary Table 3.** Residues conserved in all P4-ATPases (Conservation color codes: green – highly conserved, light green – loosely conserved, white – not conserved)

**Supplementary Table 4.** Residues conserved in all P-type ATPases (Conservation color codes: green – highly conserved, light green – loosely conserved, white – not conserved)

**Supplementary Table 5.** Residues conserved in P4-ATPases but diverged between P4A/C and P4B-ATPases (Conservation color codes: green – highly conserved, light green – loosely conserved, white – not conserved)

**Supplementary Table 6.** Residence time ranges of the characterized lipids with transmembrane helices and linkers of Neo1. All listed timescales are in μs. The total simulation time was 6 μs.

**Supplementary Table 7.** The disease mutations that have been characterized in various P4-ATPases, along with their phenotypes and biochemical effects, if characterized. The residue names and numbers for Neo1, Drs2 and ATP9A have been provided for comparison with the figures.

