## Supplementary Tables for "Structural Determinants for the Monomeric and Phospholipid-Transport P4B-ATPase via comprehensive *in silico* analysis of P4-ATPase structures": Table_S6.docx

| **TM helix** | **Range (μs)** | | | | | | | |
| --- | --- | --- | --- | --- | --- | --- | --- | --- |
|  | **POPE** | | **CHOL** | | **POPI** | | **POPS** | |
|  | **Min** | **Max** | **Min** | **Max** | **Min** | **Max** | **Min** | **Max** |
| **TM1** | 0 | 0.306 | 0 | 5.723 | 0 | 1.66 | 0 | 0.09 |
| **TM1-2 linker** | 0 | 0.132 | 0 | 0.026 | 0 | 0.62 | 0 | 0.081 |
| **TM2** | 0 | 0.236 | 0 | 2.775 | 0 | 3.541 | 0 | 0.148 |
| **TM3** | 0 | 0.093 | 0 | 0.171 | 0 | 0.367 | 0 | 0.081 |
| **TM3-4 linker** | 0 | 0.057 | 0 | 0.022 | 0 | 0.058 | 0 | 0.035 |
| **TM4** | 0 | 0.714 | 0 | 0.616 | 0 | 1.734 | 0 | 0.07 |
| **TM5** | 0 | 0.047 | 0 | 0.042 | 0 | 0.218 | 0 | 0.052 |
| **TM5-6 linker** | 0 | 0.015 | 0 | 0 | 0 | 0.052 | 0 | 0.001 |
| **TM6** | 0 | 0.171 | 0 | 0.134 | 0 | 0.572 | 0 | 0.056 |
| **TM7** | 0 | 0.418 | 0 | 0.786 | 0 | 1.109 | 0 | 0.492 |
| **TM7-8 linker** | 0 | 0.021 | 0 | 0 | 0 | 0.034 | 0 | 0.021 |
| **TM8** | 0 | 0.198 | 0 | 0.581 | 0 | 0.547 | 0 | 0.001 |
| **TM8-9 linker** | 0 | 0.255 | 0 | 0.057 | 0 | 0.723 | 0 | 0.08 |
| **TM9** | 0 | 0.179 | 0 | 0.532 | 0 | 0.369 | 0 | 0.042 |
| **TM9-10 linker** | 0 | 0.255 | 0 | 0 | 0 | 0.278 | 0 | 0 |
| **TM10** | 0 | 0.189 | 0 | 0.233 | 0 | 0.706 | 0 | 0.463 |
