## Supplementary Tables for "Structural Determinants for the Monomeric and Phospholipid-Transport P4B-ATPase via comprehensive *in silico* analysis of P4-ATPase structures": Table_S7.docx

| **Drs2** | **Neo1** | **ATP9A** | **Protein** | **Mutant** | **TM helix** | **Disease** | **Characterized biochemical effects** | **References** |
| --- | --- | --- | --- | --- | --- | --- | --- | --- |
| Q220 | E192 | N75 | ATP8B1 | E114Q | TM1 | Intrahepatic Cholestasis of Pregnancy |  | (Dixon et al. 2017) |
| T233 | V205 | L88 | ATP8B1 | L127P, L127V | TM1 | L127V - Intrahepatic cholestasis, L127P - Progressive familial intrahepatic cholestasis type 1 | Expression similar to wild-type at PM, interaction with CDC50 unaffected. Effect on substrate transport varied in different P4-ATPases (ATP8B1, ATP8A2 - PS, PE, PC transport affected; Dnf2 - PC transport unaffected) | (Narchi et al. 2017; van der Velden et al. 2010; Stone et al. 2012; Gantzel et al. 2017; Klomp et al. 2004) |
| Q237 | Q209 | Q92 | ATP11A | Q84E | TM1 | Developmental delays and neurological deterioration | GOF mutation that allowed novel PC transport ability, caused death and neurological deficits in ATP11A mouse models. PC suggested to bind the Glu residue due to theoretical positive charge of the quaternary amine at neutral pH | (Qian et al. 2026; Segawa et al. 2021) |
| E268 | D240 | E123 | ATP11A | E114G | TM2 | Developmental delays and neurological deterioration | Increased PC incorporation compared to WT | (Calianese et al. 2024) |
| L451 | L418 | L307 | ATP8B1 | I344F | TM3 | Benign recurrent intrahepatic cholestasis | Normal expression at PM (except in ATP11A). Affected substrate transport affinity and/or activity in tested flippases (ATP8A2, ATP8B1, ATP11A, Dnf2) | (Stone et al. 2012; Gantzel et al. 2017; Sun et al. 2020; Takatsu et al. 2014; Klomp et al. 2004) |
| V506 | I455 | I343 | ATP11C | I355K | TM4 | Defective B-cell development, anemia and cholestasis | Similar expression to WT, reduced PS-dependent ATPase activity | (Liou et al. 2019) |
| I508 | V457 | I345 | ATP8A2 | I376M | TM4 | Severe autosomal recessive neurological diseases (phenotypes including mental retardation, mild cerebellar and cerebral atrophy and truncal ataxia) | Abolished PS and PE transport in ATP8A2 | (Heidari et al. 2021; Coleman et al. 2014; Mogensen et al. 2024) |
| S509 | S458 | S346 | ATP8B1 | S403Y | TM4 | Progressive Familial Intrahepatic Cholestasis Type 1 | Interacts with a water molecule which is replaced by phospholipid substrate during occlusion. Reduced ATPase activity in the presence of PS and the activator taurocholic acid | (Klomp et al. 2004; Dieudonné et al. 2023; Cheng et al. 2022) |
| E1042 | Q970 | Q873 | ATP8B1 | E981K | TM6 | Progressive Familial Intrahepatic Cholestasis Type 1 | Decreased expression and substrate transport in biochemical studies. Suggested to work with a proximal residue to control substrate entry. Characterization of the same mutant in Dnf2 had contradictory results with expression unaffected but decreased Lem3 interaction; Increased affinity and decreased substrate transport rate | (Hayashi et al. 2018; Stone et al. 2012; Gantzel et al. 2017) |
| N1050 | A978 | S881 | ATP8A2 | N917D | TM6 | Cerebellar ataxia, mental retardation, and disequilibrium 4 (CAMRQ4) | N917A - PS translocation abolished. Mutant blocks dephosphorylation. Identified as the most critical residue of TM5-6 in an alanine scanning mutagenesis study. N917A/D/E/H/L/Q/R mutants all exhibited very low activity | (Alsahli et al. 2018; Mikkelsen et al. 2019; Tadini-Buoninsegni et al. 2023) |
| L1073 | T1000 | M904 | ATP8B1 | S1012I | TM6 | Childhood-Onset Progressive Intrahepatic Cholestasis | Hyperbilirubinemia | (Deng et al. 2012) |
| G1101 | S1028 | S931 | ATP8B1 | G1040R | TM7 | Progressive Familial Intrahepatic Cholestasis Type 1 | ATP8B1 - reduced expression at plasma membrane; - Dnf2 G1320R - Expression comparable to wild-type, protein unable to transport phospholipids, unable to interact with Lem3 | (Stone et al. 2012; Takatsu et al. 2014) |
